# Lifestyle activities in mid-life contribute to cognitive reserve in late-life, independent of education, occupation and late-life activities

**DOI:** 10.1101/267831

**Authors:** D Chan, M Shafto, RA Kievit, FE Matthews, M Spinks, M Valenzuela, Cam-CAN, RN Henson

## Abstract

**Introduction:** This study tested the hypothesis that mid-life intellectual, physical and social activities contribute to cognitive reserve (CR).

**Methods:** 205 individuals (196 with MRI) aged 66-88 from the Cambridge Centre for Ageing and Neuroscience (www.cam-can.com) were studied, with cognitive ability and structural brain health measured as fluid IQ and total grey matter volume, respectively. Mid-life activities were measured using the Lifetime of Experiences Questionnaire.

**Results:** Multivariable linear regression found that mid-life activities (MA) made a unique contribution to late-life cognitive ability independent of education, occupation and late-life activities. Crucially, MA moderated the relationship between late-life cognitive ability and brain structure, with the cognitive ability of people with higher MA less dependent on their brain structure, consistent with the concept of CR.

Conclusions. Mid-life intellectual, physical and social activities contribute uniquely to CR. The modifiability of these activities has implications for public health initiatives aimed at dementia prevention.

## 1. Introduction

### Participants, materials and analyses

The concept of cognitive reserve is used to explain why some individuals maintain cognitive ability despite impaired brain health as a consequence of ageing and diseases such as Alzheimer’s disease (Stern 2012). Determination of contributors to cognitive reserve is therefore important for “successful” ageing and prevention of dementia. While epidemiological evidence suggests that education and occupation contribute to cognitive reserve (Richards and Deary 2005), there is increasing interest in the additional contribution of other activities undertaken in midlife (MAs), given their potential modifiability. This interest is amplified by evidence that mid-life activities of a social or intellectual nature are associated with higher late-life cognitive ability, after adjusting for childhood cognitive ability (Gow et al. 2017), and by a recent review concluding that low levels of physical and social activity in adulthood represent key risk factors for dementia (Livingston et al. 2017).

Rigorous definitions of cognitive reserve not only predict that lifestyle factors will relate to late-life cognitive ability, but also that such factors will moderate the relationship between cognitive ability and brain structure. Specifically, the cognitive ability of individuals with high cognitive reserve should be less dependent on brain structure than those with low cognitive reserve, possibly as a result of compensatory functional network reorganisation (Stern 2017). The present study therefore asked whether MAs contribute to cognitive reserve by testing two hypotheses: i) MAs contribute to late-life cognitive ability independent of early-life education, mid-life occupation and late-life activities, and ii) MAs moderate the relationship between cognitive ability and brain structure, such that the relationship is weaker in people with higher MA.

Two hundred and five individuals (93 female) aged 66-88 years were selected from the Cambridge Centre for Ageing and Neuroscience (Cam-CAN, www.cam-can.com, Shafto et al 2014) cohort. Cognitive ability was measured using the Cattell Culture Fair test of fluid intelligence (1971) and lifestyle activities by the Lifetime of Experiences Questionnaire (LEQ, Valenzuela and Sachdev 2007), modified for UK participants.

The LEQ measures a broad range of cognitively-stimulating experiences and activities during three life phases: youth, 13-29 years; mid-life, 30-64 years; and late-life, 65 years onwards. Within each phase, further details about activities *specific* to that time of life (e.g education in youth) were solicited, as well as *nonspecific* activities applicable to any phase (e.g socialising). The youth specific score (YS, or education) was derived from the UK’s National Career Service categories, multiplied by number of years at each category. The mid-life specific score (MS, or occupation) was based on the standard occupational classification codes from the UK Office of National Statistics, summed across seven mid-life periods. The late-life specific score (LS, or post-retirement activities) reflected social and intellectual activities such as travel or participation in volunteer organisations. Scores for non-specific activities in youth (YA), mid-life (MA) and late-life (LA) were summed across seven questions about social, intellectual and physical activities. These addressed participation in i) travel, ii) social outings, iii) playing a musical instrument, iv) artistic pastimes, v) physical activity (mild, moderate, vigorous), vi) reading, vii) speaking a second language. Each of the six resulting scores (two types of activity, specific and nonspecific, across the three life phases) was scaled to a score from 0-10.

T1- and T2-weighted 1mm isotropic MRI scans were available for 196 participants. Brain structure was measured in terms of total grey matter volume (TGM, see Taylor et al. 2017). Two participants with outlying adjusted TGM values were removed.

Analysis used linear regression after all variables were standardized. Data and analysis scripts are available here: http://www.mrc-cbu.cam.ac.uk/people/rik.henson/personal/data4papers

## 2. Results

The covariance and correlation of Cattell, LEQ scores and Age are shown in Supplementary Table 1. All LEQ scores were significantly positively correlated with each other, and with Cattell.

Multivariable regression of late-life cognitive ability on the LEQ scores, with age and sex as covariates, showed a strong overall association (adjusted R^2^=0.355, F(8,196)=15.0, p<.001). In addition to the expected negative effect of age, the normalized coefficients in Table 1 revealed a unique, positive contribution of YS (i.e, education), replicating previous studies such as Richards and Deary (2005). More interestingly, MA also made an independent positive contribution after adjustment for all other factors, with the largest effect size. No other LEQ-based category, including the current late-life activities being performed by the individuals (reflected in both LS and LA), made an independent contribution.

**Table 1.** Results of multivariable linear regression of late-life cognitive ability (Cattell) against six lifetime experience scores from the LEQ, plus age and sex.

|  | Normalised Coefficient | Standard Error | p-value (df=196) |
| --- | --- | --- | --- |
| Young specific activities | +0.259 | 0.071 | **<0.0001** |
| Young nonspecific activities | +0.028 | 0.078 | 0.723 |
| Midlife specific activities | +0.096 | 0.069 | 0.164 |
| Midlife nonspecific activities | +0.324 | 0.075 | **<0.0001** |
| Late life specific activities | +0.097 | 0.062 | 0.110 |
| Late life nonspecific activities | -0.098 | 0.075 | 0.195 |
| Age | -0.287 | 0.058 | **<0.0001** |
| Sex | -0.082 | 0.064 | 0.201 |

To examine whether MAs contributed not just to late-life cognitive ability but also to cognitive reserve, rigorously defined, we determined whether MAs moderated the relationship between late-life cognitive ability and brain structure (as indexed by TGM).

We first adjusted Cattell and TGM scores for 1) Education (YS score), 2) Total Intracranial Volume (TIV), 3) Age and 4) Sex. Linear regression showed an interaction between TGM and MA in predicting Cattell (normalized interaction coefficient = −0.722, standard error = 0.331, p=0.030). This moderating term was negative, meaning that the relationship between cognitive ability and brain health diminished with higher MA, as predicted by cognitive reserve theory. This effect is visualized in Figure 1, where a median split is used to divide participants into low and high MA groups: a more positive slope can be seen in the low MA group than in the high MA group.

**Figure 1.**
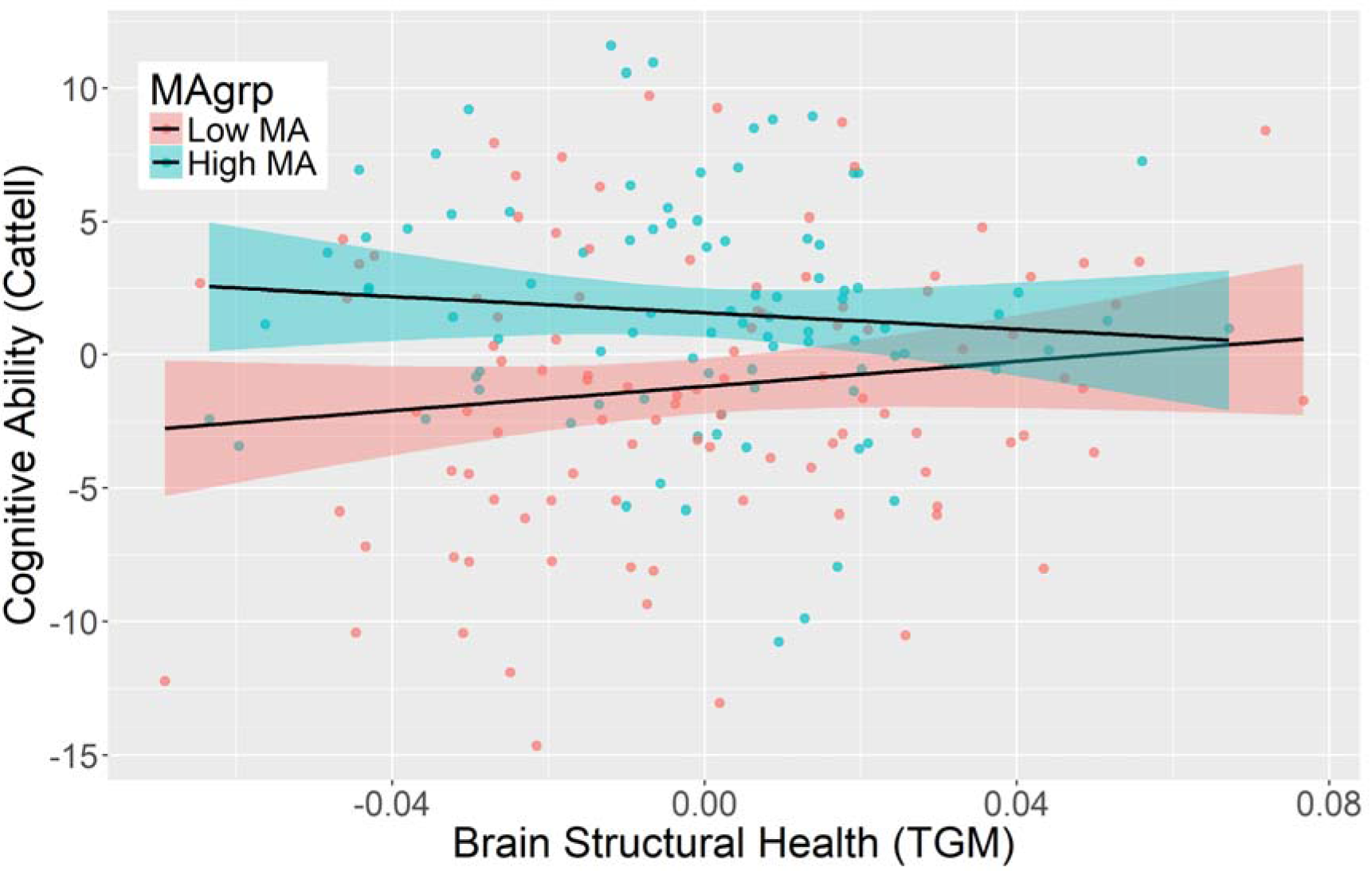
Relationships between cognitive ability and brain structure in the high and low MA groups. Both variates were adjusted for youth-specific activities (education), total intracranial volume, age and sex.

## 3. Discussion

This study tested the hypothesis that lifestyle activities in mid-life contribute to the cognitive reserve that supports cognitive ability in late-life. Consistent with this, we found that general mid-life activities (MA) make an independent contribution to late-life cognition over and above age, sex, education, occupation and current (late-life) activities, replicating the findings of Gow et al (2017). However, a rigorous determination of cognitive reserve requires further evidence that it moderates the association between a brain state and cognitive outcome (e.g., Brayne et al 2010). Importantly, we found such a moderation by MA on the relationship between total grey matter volume and cognitive ability, such that cognitive ability in those older individuals who had been involved in rich and varied lifestyle activities in mid-life were less dependent on their current structural brain health.

This evidence that midlife activities contribute to cognitive reserve may have ramifications for the primary prevention of dementia. These activities reflect lifestyle choices, and are therefore amenable to modification. Our observation that the impact of MA was independent of educational attainment and occupational status, suggests that a public health initiative aimed at boosting cognitive reserve via enhancement of MA is generalizable to the entire adult population. The importance of such initiatives for primary prevention of dementia is underscored by the total failure to date to identify interventions for secondary prevention of dementia.

This study does not address the relative contribution to cognitive reserve of the various intellectual, social and physical components of the LEQ midlife activities score. Many studies have suggested that physical activity reduces future dementia risk (Richards et al 2003, Middleton et al 2010), but the UK Whitehall II study found no association between physical activity and subsequent 15 year cognitive decline (Sabia et al 2017). Similarly, Gow et al (2017) found that mid-life intellectual and social activities, but not physical activity, were associated with late-life cognitive health. Other studies have also highlighted the benefits of intellectual and social activity (Akbaraly et al. 2009, Köhncke et al 2016). Future work, complementing the LEQ with specific measures for each of these MA components, will be needed to clarify this issue.

Another key objective for future research is identification of the neural mechanisms by which MA supports late life cognitive ability. One possibility is that MA alters the functional networks subserving cognitive processes, rendering them more resilient to structural brain atrophy (Marques et al. 2016, Stern 2017). Identification of the underlying mechanisms will have relevance not just for the understanding of cognitive reserve but also for the emerging concept of brain maintenance (Nyberg et al. 2012).

There are important limitations of this study. First, LEQ measures are self-report, raising the possibility that more cognitively healthy older people remember more lifetime activities. Second, a limitation of cross-sectional studies is the possibility of reverse causation, namely that a higher cognitive ability throughout life triggers the pursuit of more beneficial lifestyle activities, rather than vice versa. Third, LEQ scores were highly positively correlated, so we cannot infer that other variables (e.g, occupation) play no role in late-life cognitive ability. These limitations could be addressed by intervention studies targeting specific aspects of mid-life activity, with later life cognitive ability as the outcome measure, though such studies would require a timeframe of 20-30 years.

In conclusion, our findings suggest that lifestyle activities in mid-life can contribute to cognitive reserve and support late-life cognition. The potential modifiability of these activities has important implications for public health initiatives aimed at reducing the risk of dementia.

## Acknowledgments

DC is funded by the Cambridge National Institute for Health Research (NIHR) Biomedical Research Centre. FM is part-funded by Medical Research Council (MC U105292687). MV is funded by a National Health and Medical Research Council of Australia Career Development Fellowship. The Cambridge Centre for Ageing and Neuroscience (Cam-CAN) research was supported by the Biotechnology and Biological Sciences Research Council (grant number BB/H008217/1). Additional support to RH and RK was from the Medical Research Council (grants SUAG/010 RG91365 and SUAG/014 RG91365) and the European Union’s Horizon 2020 research and innovation programme under grant agreement No 732592. We are grateful to the Cam-CAN respondents and their primary care teams in Cambridge for their participation in this study. The Cam-CAN corporate author consists of the people named here: http://www.cam-can.com/index.php?content=corpauth#12.

---

**Supplementary Table 1.** Variance (diagonal), covariance (upper triangle) and correlation (lower triangle) of Cattell, LEQ scores and Age. Last Column shows min/max range. YS = youth specific activities, YA = nonspecific youth activities, MS = mid-life specific activities; MA = nonspecific mid-life activities, LS = late-life specific activities; LA = nonspecific late-life activities. *p < 0.05.

|  | Cattell | YS | YA | MS | MA | LS | LA | Age |
| --- | --- | --- | --- | --- | --- | --- | --- | --- |
| Cattell | 37.2 | 8.88 | 4.12 | 5.98 | 4.76 | 1.99 | 2.74 | -12.2 |
| YS | +.43* | 11.5 | 3.12 | 4.24 | 1.84 | 1.03 | 1.94 | -1.78 |
| YN | +.32* | +.44* | 4.57 | 0.94 | 2.41 | 1.09 | 1.99 | -0.69 |
| MS | +.36* | +.46* | +.16* | 7.27 | 0.80 | 0.80 | 0.76 | -3.78 |
| MN | +.39* | +.27* | +.56* | +.15* | 4.00 | 0.86 | 2.04 | -0.48 |
| LS | +.18* | +.17* | +.29* | +.17* | +.24* | 3.14 | 0.97 | 0.59 |
| LN | +.26* | +.33* | +.54* | +.16* | +.59* | +.32* | 2.99 | -0.96 |
| Age | -.33* | -.09 | -.05 | -.24* | -.04 | .06 | -.09 | 34.8 |

## References

Stern Y (2012) Cognitive reserve in ageing and Alzheimer’s disease. Lancet Neurol 11, 1006–1012

Richards M, Deary IJ (2005) A life course approach to cognitive reserve: a model for cognitive aging and development? Ann Neurol. 58, 617–22.

Gow AJ, Pattie A, Deary IJ (2017) Lifecourse Activity Participation From Early, Mid, and Later Adulthood as Determinants of Cognitive Aging: The Lothian Birth Cohort 1921. J Gerontol B Psychol Sci Soc Sci. 72, 25–37

Livingston G, Sommerlad A, Orgeta V, et al. (2017) Dementia prevention, intervention, and care. Lancet. 2017 Jul 19. pii: S0140-6736(17)31363-6. doi: 10.1016/S0140-6736(17)31363-6.

Stern Y (2017) An approach to studying the neural correlates of reserve. Brain Imaging Behav 11,410–416

Shafto MA, Tyler LK, Dixon M et al. (2014) The Cambridge Centre for Ageing and Neuroscience (Cam-CAN) study protocol: a cross-sectional, lifespan, multidisciplinary examination of healthy cognitive ageing. BMC Neurol. 2014 Oct 14;14:204. doi: 10.1186/s12883-014-0204-1.

Cattell, RB (1971) Abilities: their Structure, Growth, and Action (Houghton-Mifflin)

Valenzuela MJ, Sachdev P (2007) Assessment of complex mental activity across the lifespan: development of the Lifetime of Experiences Questionnaire (LEQ). Psychol Med 37,1015–25.

Taylor, J.R., Williams, N., Cusack, R. et al (2017). The Cambridge Centre for Ageing and Neuroscience (Cam-CAN) data repository: Structural and functional MRI, MEG, and cognitive data from a cross-sectional adult lifespan sample. NeuroImage 144, 262–269

EClipSE Collaborative Members, Brayne C, Ince PG, et al (2010) Education, the brain and dementia: neuroprotection or compensation? Brain 133, 2210–6.

Richards M, Hardy R, Wadsworth ME (2003) Does active leisure protect cognition? Evidence from a national birth cohort. Soc Sci Med 56, 785–92

Middleton LE, Barnes DE, Lui LY et al. (2010) Physical activity over the life course and its association with cognitive ability and impairment in old age. J Am Geriatr Soc. 58, 1322–6

Akbaraly TN, Portet F, Fustinoni S, et al. (2009) Leisure activities and the risk of dementia in the elderly: results from the Three-City Study. Neurology 73, 854–61.

Köhncke Y, Laukka EJ, Brehmer Y, et al. (2016) Three-year changes in leisure activities are associated with concurrent changes in white matter microstructure and perceptual speed in individuals aged 80 years and older. Neurobiol Aging 41, 173–186.

Marques P, Moreira P, Magalhães R et al. (2016) The functional connectome of cognitive reserve. Hum Brain Mapp 37, 3310–22

Nyberg L, Lövdén M, Riklund K, et al (2012) Memory aging and brain maintenance. Trends Cogn Sci 16, 292–305

